# Neural dynamics at rest associated with patterns of ongoing thought

**DOI:** 10.1101/454371

**Authors:** Theodoros Karapanagiotidis, Diego Vidaurre, Andrew J. Quinn, Deniz Vatansever, Giulia L. Poerio, Elizabeth Jefferies, Daniel S. Margulies, Thomas E. Nichols, Mark W. Woolrich, Jonathan Smallwood

**Affiliations:** Department of Psychology, York Neuroimaging Centre, University of York, York YO10 5DD, United Kingdom; Oxford Centre for Human Brain Activity, Wellcome Centre for Integrative Neuroimaging, Department of Psychiatry, University of Oxford, Oxford OX3 7JX, United Kingdom; Department of Psychology, University of Sheffield, Sheffield S1 2LT, United Kingdom; Brain & Spine Institute (ICM), National Center for Scientific Research, Paris 75013, France; Oxford Centre for Functional MRI of the Brain, Wellcome Centre for Integrative Neuroimaging, Nuffield Department of Clinical Neurosciences, University of Oxford, Oxford OX3 9DU, United Kingdom

## Abstract

Conscious experience is dynamic, and its fluidity is particularly marked when attention is not occupied by events in the external world and our minds are free to wander. Our study used measures of neural function, and advanced analyses techniques to examine how unconstrained neural state transitions relate to patterns of ongoing experience. Neural activity was recorded during wakeful rest using functional magnetic resonance imaging and Hidden Markov modelling identified recurrent patterns of brain activity constituting functional dynamic brain states. Individuals making more frequent transitions between states subsequently described experiences highlighting problem solving and lacking unpleasant intrusive features. Frequent switching between states also predicted better health and well-being as assessed by questionnaire. These data provide evidence that the fluidity with which individuals shift through dynamic neural states has an impact on the nature of ongoing thought, and suggest that greater flexibility at rest is an important indicator of a healthy mind.

## Introduction

William James (James, 1890) emphasised experience unfolds dynamically over time, using the analogy of a “stream of consciousness”. The fluidity of experience is clearly illustrated by the fact that our attention tends to flit from topic to topic, particularly when we are not focused on events in the external world (Smallwood and Schooler, 2006, 2015). Such “mind-wandering” is a broad class of experience (Seli et al., 2018) that is common in daily life (Killingsworth and Gilbert, 2010), consistent across cultures (Singer and McCraven, 1961) and declines with age (Giambra, 1989). Mind-wandering is linked with improved creativity (Baird et al., 2012) and problem solving (Medea et al., 2016) suggesting that it may facilitate our ability to navigate the complex social environment in which we exist (Stawarczyk et al., 2011; Smallwood and Schooler, 2015). In contrast, less flexible patterns of conscious thought can get “stuck” in a cycle of intrusive rumination, with detrimental consequences. Unpleasant moods, for example, can encourage patterns of negative thoughts about the past (Smallwood and O’Connor, 2011; Poerio et al., 2013) which in turn reduce subsequent mood (Ruby et al., 2013). These studies highlight links with mental health since patterns of recurrent rumination can maintain states of anxiety and depression (Watkins, 2008). It is apparent that mind-wandering has both beneficial and detrimental associations and what dissociates these two extremes may be the flexibility with which cognition dynamically unfolds over time.

Recent advances in neuroimaging analysis make it possible to test the impact that covert cycling through neural states has on patterns of ongoing experience. Traditional functional connectivity analyses exploit temporal correlations in brain activity to identify the spatial extent of large-scale distributed neural networks (Smith et al., 2013b). These networks are relevant to cognition given their associations with population variation in intelligence and well-being (Smith et al., 2015; Finn et al., 2015), as well as more specific cognitive factors such as cognitive flexibility (Vatansever et al., 2017) and creativity (Beaty et al., 2014). These traditional methods, however, describe neural networks in a static (time-averaged) manner, and so do not explicitly address the temporal profile of neural function. Contemporary research has begun to address the dynamic properties of neural activity in a more direct manner using Hidden Markov Modelling (HMM). This method identifies ‘states’, defined as recurrent patterns of neural activity, and studies have shown that states revealed in this manner track cognitive processes during tasks (Gonzalez-Castillo et al., 2015), and at rest, relate neural processing to cognitive flexibility, life satisfaction, anger and perceived stress (Vidaurre et al., 2017b). By providing a quantified description of covert states and how individuals transition between them, HMM allows us to test how the underlying neural dynamics impact upon aspects of ongoing experience.

The current study mapped patterns of intrinsic neural activity using functional Magnetic Resonance Imaging (fMRI) in a large cohort of individuals (N = 169) while they rested in the scanner. We interrogated undisturbed wakeful rest, rather than probing during a task (e.g. (Sormaz et al., 2018)) for two reasons. First, wakeful rest is conducive to experiences such as mind-wandering (Smallwood et al., 2009). Second, the absence of external interruptions during rest ensures that neural dynamics unfold in a relatively natural way. At the end of the scan, participants retrospectively described their experience, answering a set of questions based on those used in prior studies exploring variability in static functional connectivity across subjects (Smallwood et al., 2016; Karapanagiotidis et al., 2017). In a separate session, participants completed a set of validated measures of physical and mental health. Following the HMM inference of the dynamic states, we calculated indices of the temporal flexibility of neural function and the time spent in a particular state. We used these metrics as explanatory variables in separate regression analyses examining their relationship to descriptions of subjective experience reported at the end of the scan, as well as their links to the validated measures of health and well-being. To foreshadow our results, individuals who made more transitions between states described experiences emphasising problem solving and fewer intrusive thoughts. They also scored higher on measures of health and well-being. These data provide evidence that the fluidity with which an individual shifts through unconstrained states is linked to the emergence of more apparently useful patterns of ongoing thought.

## Results

### Describing experience and well-being

In order to identify the measures of experience with the best reliability, we repeated the resting state scanning session for a subset of our sample (N = 40) approximately 6 months later. This revealed 6 questions that were consistent across sessions (Fig. 1a): normal thoughts (i.e. experiences that I often have, intraclass correlation coefficient = 0.28, p = 0.035), deliberate vs spontaneous thoughts (ICC = 0.27, p = 0.044), intrusive thoughts (ICC = 0.29, p = 0.034), problem-solving thoughts (ICC = 0.454, p = 0.001), thoughts about the here and now (ICC = 0.281, p = 0.036) and thinking in images (ICC = 0.3, p = 0.026). We used these items in subsequent analyses.

**Figure 1.**
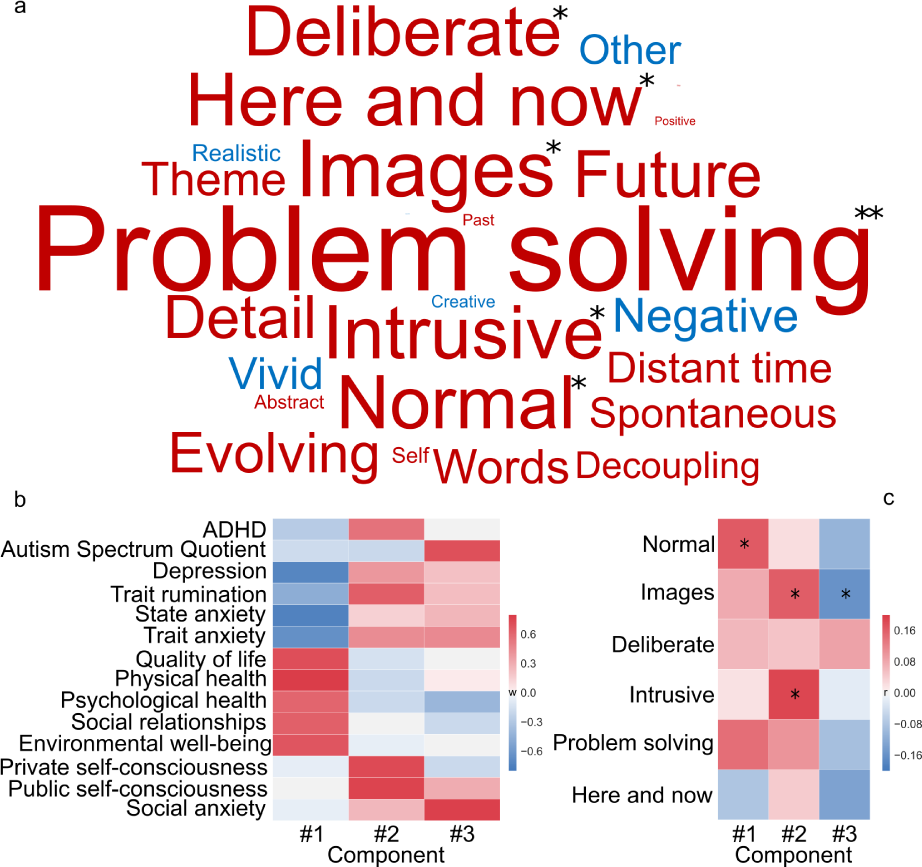
Behavioural variables. **a)** Test-retest reliability of responses to items at the end of two resting state fMRI scans for 40 participants. Intraclass correlation coefficient (ICC) visualised using a word cloud. Font size represents the absolute ICC value and font colour its sign (red for positive and blue for negative values). * p < 0.05, ** p = 0.001. **b**) Component weights from a principal component analysis on the participants’ responses to physical and mental health questionnaires. **c**) Correlation between the 6 thought probes that were significantly reliable in (a) and the component scores from (b). Analyses controlled for age, gender and motion during the resting state fMRI scan.

To reduce the dimensional structure of the measures of physical and mental health, we performed a principal components analysis decomposition with varimax rotation. This identified three principal components with eigenvalues greater than 1 (Fig. 1b). Component 1 loaded positively on measures of physical health and psychological well-being and negatively on indices of depression and anxiety. Component 2 loaded on self-consciousness, ADHD and rumination. Component 3 loaded on social anxiety and autism.

Exploring links between measures of well-being and items describing experience at rest, we found a positive correlation between normal patterns of thoughts with component 1 (r = 0.17, p < 0.05), intrusive thoughts (r = 0.19, p < 0.05) and thinking in images (r = 0.16, p < 0.05) with component 2 and a negative correlation of thinking in images (r = −0.16, p < 0.05) with component 3 (Fig. 1c). However, none of these associations were significant after controlling for multiple comparisons.

### Describing neural dynamics

Recent work has highlighted that patterns of ongoing thought can be meaningfully related to interactions between multiple large-scale networks (Golchert et al., 2017; Hasenkamp et al., 2012; Mooneyham et al., 2017). To capture network-to-network relationships, we conducted spatial independent component analysis (ICA) on the resting state fMRI data, identifying 15 functional networks and then ran the HMM (variational Bayes) inference on their time courses (Vidaurre et al., 2017b). HMM allows the whole multi-dimensional fMRI time course to be decomposed into a set of recurring functional states, each characterised by unique patterns of network activity. In determining the number of states for the HMM, we tested models with 7, 9 and 12 states and evaluated the stability of the decompositions by running the algorithm 100 times in each case. Models with fewer states were found to be generally more stable (Fig. S3) hence we discuss the 7 state solution in the main body of our paper. Out of the 100 runs of the algorithm, we selected the best solution (optimal HMM) according to the free energy (Quinn et al., 2018), a measure related to the Bayesian inference process that ranks the solutions based on their complexity and accuracy in representing the data (see Methods). All 7 states as inferred by the optimal HMM are presented in Figure S2. Our results utilise dynamic metrics inferred from this optimal HMM solution, however they were also robust when taking into account the run-to-run variability across all 100 models produced by the algorithm (Vidaurre et al., 2018) (see SI Results).

From the HMM decomposition, a number of summary metrics describing the dynamics in the data can be extracted. The overall switching rate across states (histogram shown in Fig. 2a), represents the lability of the dynamics. The transition probabilities across states (see Fig. 2b) describe how likely transitions between different states are. Several brain states have a high probability of transitioning into State 4 (*µ*_*tS*4_ = 0.22) and, to a lesser degree, to State 3 (*µ*_*tS*3_ = 0.17, the full asymmetric transition probability matrix is presented in Fig. S4). Finally, the fractional occupancy (FO) describes the proportion of time spent in each state. Most states had comparable FO with the exception of State 5, which was lower than all other states (Fig. 2c).

**Figure 2.**
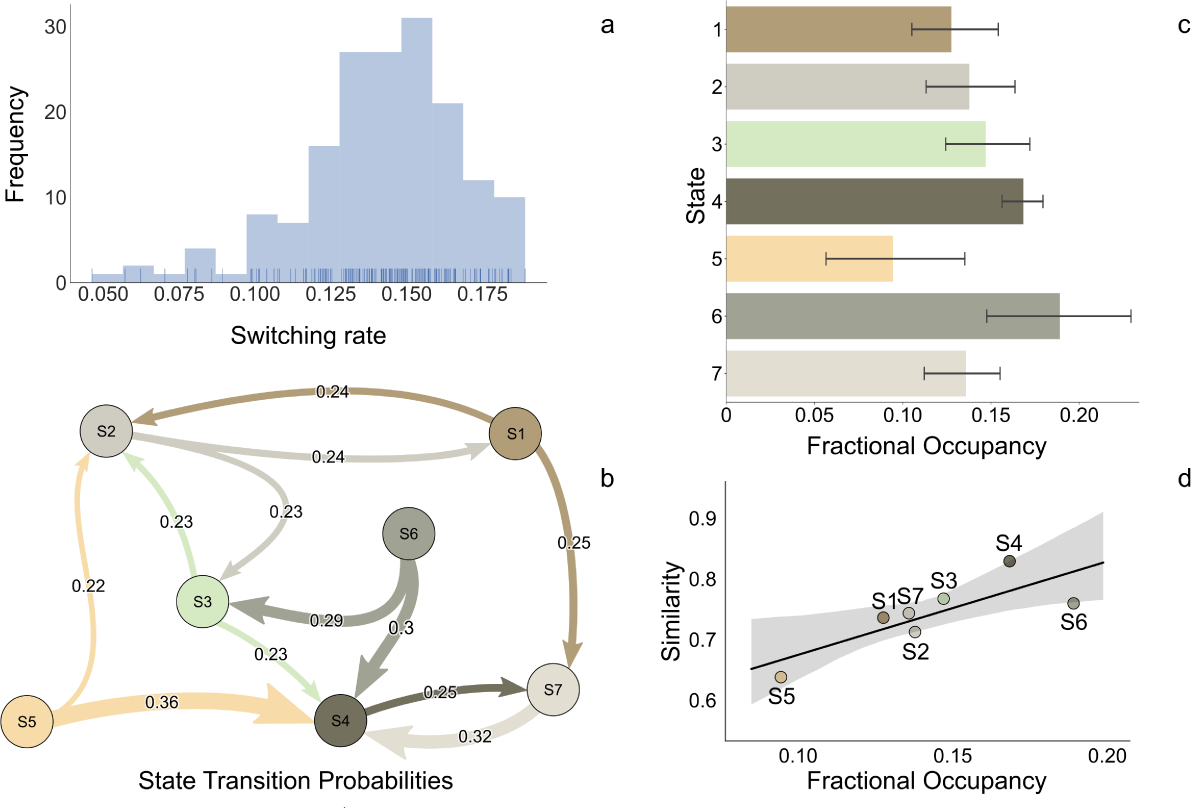
Dynamic metrics. **a**) Histogram of subjects’ switching rate between all states. **b**) The states’ transition probabilities as a thresholded graph, with edge width weighted by the respective probability (the probabilities of staying in the same state are not shown). **c**) Bar plots of the average fractional occupancy of each state over subjects (±99.9% bootstrap confidence intervals). **d**) Scatter plot of the average fractional occupancy of each dynamic state and its similarity with the mean static functional connectivity across subjects.

As expected, relating the correlation matrix of each state with the group static functional connectivity (i.e. the time averaged partial correlation matrix), we found that the more time spent in a particular dynamic state, the more similar states were (in terms of whole-brain connectivity) to the average static functional connectivity pattern (p = 0.031) (Fig. 2d). This is consistent with the idea that the patterns of static connectivity observed across the entire period of rest can be understood as the superposition of the dynamic states identified through the HMM.

### Relationship between brain dynamics and ongoing experience

Having identified metrics that describe dynamic brain activity at rest, we tested whether these relate to descriptions of subjective experience at the end of the scan.

#### Overall dynamics and ongoing experience

We used overall state switching rate during the resting state scan as an explanatory variable in a multivariate GLM, combined with permutation testing, to determine statistical significance. We identified a significant association with concurrent thoughts at the end of the scan (p = 0.003) (Fig. 3, top). Follow-up univariate analyses and permutation testing showed that increased switching between states was predictive of less intrusive thoughts (r = −0.21, p = 0.014 uncorrected, p = 0.04 FDR-corrected for 6 tests (number of questions)), and more related to problem solving (r = 0.20, p = 0.005 uncorrected, p = 0.03 FDR-corrected) (Fig. 3, bottom).

**Figure 3.**
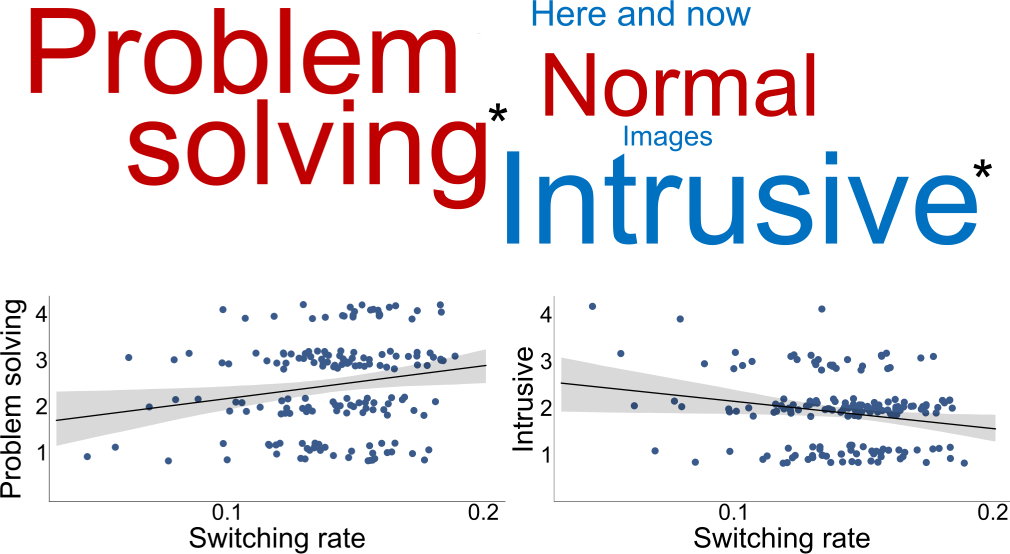
Overall dynamics associated with ongoing experience. Correlation coefficients between thoughts and state switching rate visualised using a word cloud (top) and the associated scatter plots for the significant relationships (bottom). Font size in world cloud represents the absolute correlation value and font colour its sign (red for positive and blue for negative correlations). * p < 0.05 (FDR-corrected).

Parallel analyses indicated similar associations with the switching rate for an HMM decomposition of 9 states (p = 0.019) and 12 states (p = 0.0005) (SI Results), demonstrating this relationship was robust to the number of *a priori* states selected. No other questionnaire items showed a significant relationship with switching rate.

#### State-specific dynamics and ongoing experience

We also explored the relationship between the FO of the states and the descriptions of ongoing experience. Using the FO of each state as an explanatory variable in separate multivariate GLM, we found a significant relationship between the FO of state 3 and the FO of state 5 with thoughts (p = 0.018 and p = 0.036 respectively, uncorrected) (Fig. 4b). State 3 was dominated by three motifs of increased connectivity between (i) the default mode (DMN) and language network, (ii) lateral and medial visual cortex and (iii) the precuneus and saliency network. Whereas, state 5 was marked by strong coupling between sensory systems (sensorimotor and auditory) (Fig. 4a). These results, however, did not pass correction for multiple testing (p = 0.13, FDR-corrected for 7 tests (number of states)), rendering them less robust than the effects of the overall dynamics and so we do not consider them in detail further.

**Figure 4.**
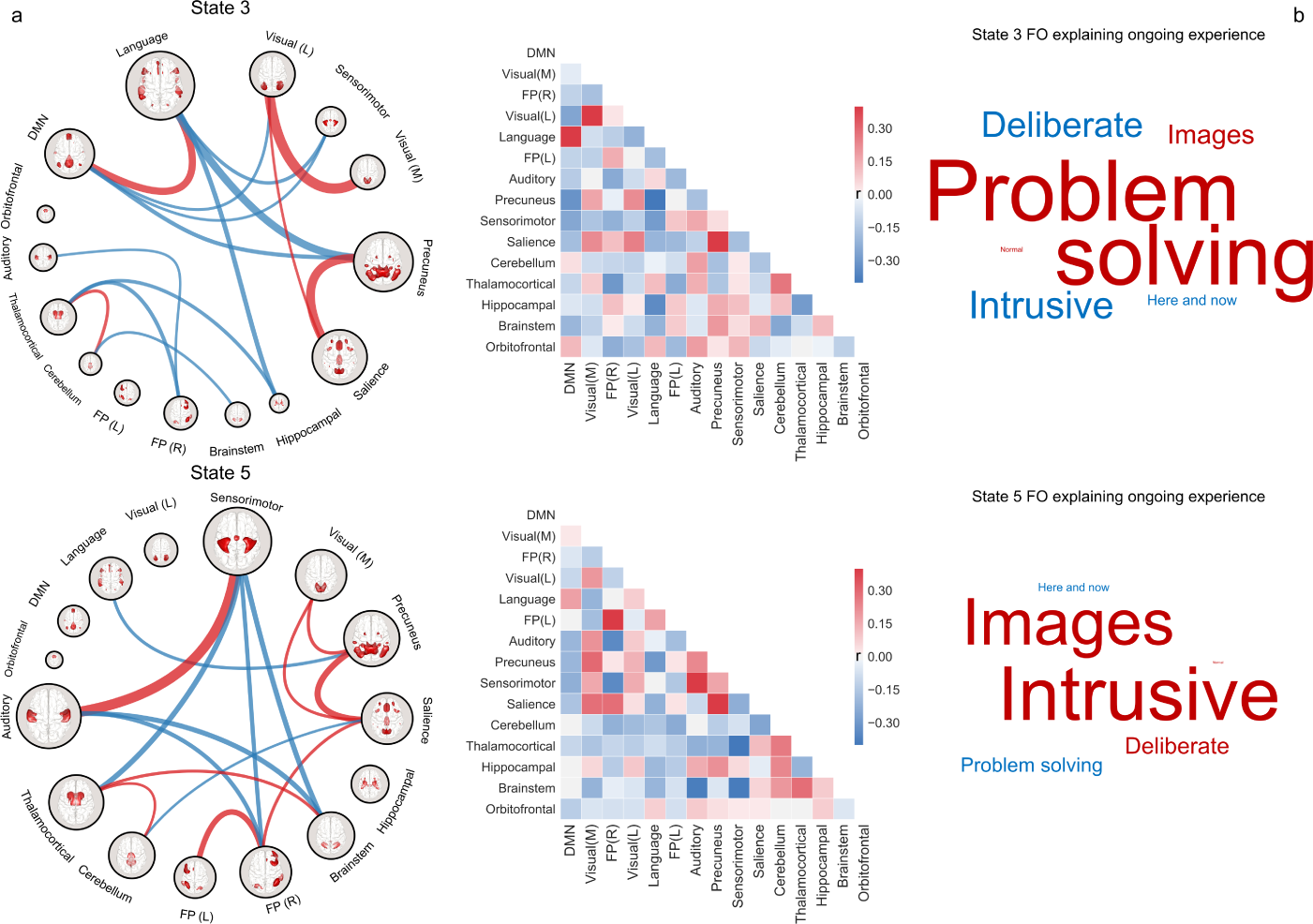
State-specific dynamics associated with ongoing experience. **a**) This panel presents chord graphs of states 3 and 5 (left) along with their corresponding correlation matrices (middle). The chord graphs were thresholded, for visualisation, at the 85*th* percentile of absolute correlation values for each state. Red lines refer to positive and blue lines refer to negative correlations, with edge width representing the magnitude of the correlation. Node size shows the node’s “strength”, computed as the sum of the absolute correlation of each network with the rest of the brain. **b**) Correlation coefficients between thoughts and fractional occupancy of state 3 (top) and state 5 (bottom) visualised using word clouds. Font size in world clouds represents the absolute correlation value and font colour its sign (red for positive and blue for negative correlations). Note, the relationship between the FO of the states and ongoing experience does not pass correction for multiple comparisons.

### Relationship between brain dynamics and measures of general health and well-being

After demonstrating links between brain dynamics and reports of experience, we next examined associations between dynamic neural metrics and measures of physical and mental health. The analysis showed that state switching rate was related to well-being (p = 0.04). In particular, increased dynamic brain activity, as identified by our temporal classification, was linked with increased general health and psychological well-being and with less depression and anxiety (Component 1 in Fig. 1b) (r = 0.20, p = 0.007 uncorrected, p = 0.02 FDR-corrected for 3 tests (number of PCA components), Fig. 5). The FO of the HMM states was not significantly correlated with any of the 3 PCA components.

**Figure 5.**
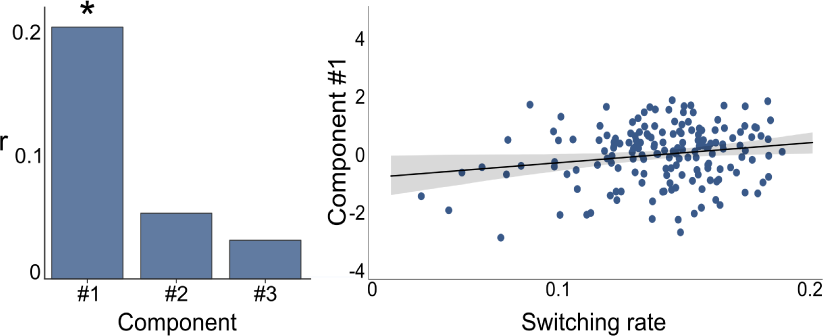
Brain dynamics associated with physical and mental health. Correlation coefficients between PCA components from the physical and mental health scores and state switching rate and the associated scatter plot for the significant relationship with component 1. * p < 0.05 (FDR-corrected).

## Discussion

Consistent with the view that temporal dynamics are an integral aspect of unconstrained cognition, we found the degree of switching between different intrinsic neural states was associated with self-reported experiential states emerging during this period. Greater flexibility was linked to less intrusive thoughts and more related to attempts at problem solving, as well as better physical and mental well-being. In constrast, we found only modest evidence relating patterns of experience to the occupancy of individual states. These data support contemporary accounts of ongoing experience that emphasise temporal dynamics as important for understanding patterns of concurrent experience (Smallwood, 2013; Christoff et al., 2016; Kucyi et al., 2016), and we consider the significance of this observation for our understanding of ongoing cognition.

Our study suggests the flexibility of intrinsic neural activity may be a mechanism determining whether patterns of ongoing thought are linked to beneficial or deleterious aspects of cognition and behaviour (Smallwood and Andrews-Hanna, 2013). More transitions between neural states at rest was linked to patterns of experience that emphasised problem solving. Autobiographical planning and problem solving are widely accepted benefits that can emerge from unconstrained thought (Smallwood and Schooler, 2015) which have been shown to help refine personal goals (Medea et al., 2016) and reduce psycho-social stress (Engert et al., 2014). In contrast, individuals for whom neural dynamics were inflexible reported experiences that were more intrusive in nature. Intrusive thinking is central to multiple psychiatric conditions, including obsessive compulsive disorder (Najmi et al., 2009), depression/rumination (Smith and Alloy, 2009) and post-traumatic stress disorder (de Silva and Marks, 1999). Critically, greater switching between states at rest was also linked to better well-being. Together, our data suggests that more flexible neural activity is linked to more beneficial associated outcomes.

More generally, our study suggests that abstract properties of neural dynamics, such as the ease of transition between states, can be psychologically relevant (Kucyi, 2017). Previous studies have focused on the consistency of the states, for example, by documenting their heritability (Vidaurre et al., 2017b). Others have examined the influence of particular states on cognition and behaviour. Cabral and colleagues (Cabral et al., 2017), for example, found higher cognitive performance for older individuals was predicted by the prevalence of a global brain state. Our study found modest evidence that particular experiences are related to states with specific neural motifs, however, we found stronger evidence that eases of transition between states is linked to experience, and health and well-being. Based on these findings, it will be important to understand the manner through which the flexibility of cycling through different neural states contributes to cognition and also whether this can change across the life cycle and with clinical conditions. Studies have shown specific sequences of neural processing support the expression, recognition and supression of mind-wandering in experienced meditators (Hasenkamp and Barsalou, 2012), and so part of the flexibility we observe may reflect a process implicated in the monitoring of ongoing thought (Schooler, 2002). More generally, understanding how state transitions are constrained by cortical architecture (Margulies et al., 2016) and influenced by the distributed activity of neuromodulators (Harris-Warrick, 2011) are important questions for future research to address.

Finally, our demonstration that HMM can reveal covert neural states relevant to patterns of ongoing thought constitutes an important methodological advance for the study of ongoing conscious experience. Although famous for highlighting dynamics, William James argued against introspection as an approach to understanding experience, likening it to trying to “turn up the gas quickly enough to see the darkness” (James, 1890). While the last two decades have seen increased interest in investigating self-generated thoughts (Callard et al., 2013), we lack a method to study them in a manner that does not disrupt their natural dynamics (Konishi and Smallwood, 2016; Kucyi, 2017). Our study shows that the HMM captures features of neural dynamics that are relevant to ongoing thought in a manner that does not rely on introspective information for their identification. The ability of the HMM to ‘parse’ neural data into underlying temporal states, without disrupting their evolution, provides a window into how cognition unfolds without the concern that the disruptive nature of experience sampling may contaminate it. We anticipate future studies will be able to apply this method to gain unprecedented access to covert changes in cognition that are a pervasive, yet poorly understood aspect of our mental lives.

## Methods and Materials

### Participants

206 healthy participants were recruited by advert from the University of York. Written consent was obtained for all participants and the study was approved by the York Neuroimaging Centre Ethics Committee. 37 participants were excluded from analyses due to technical issues during the neuroimaging data acquisition, excessive movement during the fMRI scan (mean framewise displacement > 0.3 mm and/or more than 15% of their data affected by motion) (Power et al., 2014) or not completing the whole battery of behavioural tasks, resulting in a final cohort of N = 169 (111 females, *µ_age_* = 20.1 years, *σ_age_* = 2.3).

### Behavioural methods

We sampled the participants’ experiences during the resting state fMRI scan by asking them at the end of the scan to retrospectively report their thoughts, using a series of self-report questions. These items were measured using a 4-scale Likert scale with the question order being randomised (all 25 questions are shown in Table S1). In order to assess the participants’ physical and mental health, we administered well-established surveys at a later separate session outside of the scanner. Details about each questionnaire are presented in SI Methods. Analyses controlled for age, gender and motion during the resting state fMRI scan.

### Neuroimaging methods

#### MRI data acquisition

MRI data were acquired on a GE 3 Tesla Signa Excite HDxMRI scanner, equipped with an eight-channel phased array head coil at York Neuroimaging Centre, University of York. For each participant, we acquired a sagittal isotropic 3D fast spoiled gradient-recalled echo T1-weighted structural scan (TR = 7.8 ms, TE = minimum full, flip angle = 20°, matrix = 256×256, voxel size = 1.13×1.13×1 mm, FOV = 289×289 mm^2^). Resting-state functional MRI data based on blood oxygen level-dependent contrast images with fat saturation were acquired using a gradient single-shot echo-planar imaging sequence with the following parameters TE = minimum full (≈19 ms), flip angle = 90°, matrix = 64×64, FOV = 192×192 mm^2^, voxel size = 3×3×3 mm^3^TR = 3000 ms, 60 axial slices with no gap and slice thickness of 3 mm. Scan duration was 9 minutes which allowed us to collect 180 whole-brain volumes.

#### fMRI data pre-processing

Functional MRI data pre-processing was performed using the Configurable Pipeline for the Analysis of Connectomes (C-PAC) (Craddock et al., 2013). Pre-processing steps included motion correction by volume realignment (Friston 24-Parameter Model) (Friston et al., 1996), nuisance signal regression of the 24 motion parameters calculated in the previous step plus five nuisance signals obtained by running a principal components analysis on white matter and cerebrospinal fluid signals using the CompCor approach (Behzadi et al., 2007), slice time correction, temporal filtering 0.009-0.1 Hz, spatial smoothing using a 6mm Full Width at Half Maximum of the Gaussian kernel and normalisation to MNI152 stereotactic space (2 mm isotropic) using linear and non-linear registration (boundary-based registration) (Greve and Fischl, 2009). No global signal regression was performed.

#### Group-ICA spatial maps and time series

Following pre-processing, the neuroimaging data were masked by a 20% probabilistic whole-brain grey matter mask, temporally demeaned, had variance normalisation applied (Beckmann and Smith, 2004) and were fed into the MIGP algorithm (Smith et al., 2014). The output of MIGP, which is a very close approximation to running PCA on the temporally concatenated data, was then fed into group-ICA using FSL’s MELODIC tool (Beckmann and Smith, 2004), where spatial-ICA was applied, resulting in 16 distinct group-ICA spatial maps (SI results, Fig. S1). These group spatial maps were subsequently mapped onto each subject’s pre-processed data by running the first stage in a dual-regression analysis, which produced one time series per map per participant. After removal of one artefactual component, the remaining 15-dimensional time series for each participant with *µ* = 0 and *σ* = 1 were concatenated to form a (180 x 169) x 15 matrix and used as input for subsequent analyses.

#### Hidden Markov model

To characterise the dynamics of neural activity, we applied a hidden Markov model to the concatenated time series of the ICA networks. The inference of the model parameters was based on variational Bayes and the minimisation of free energy, as implemented in the HMM-MAR toolbox (Vidaurre et al., 2016). The HMM’s inference assigns state probabilities to each time point of the time series (i.e. reflecting how likely is each time point to be explained by each state) and estimates the parameters of the states, where each state has its own model of the observed data. Each state can be represented as a multivariate Gaussian distribution (Vidaurre et al., 2017a), described by its mean and covariance. As we were primarily interested in identifying changes in functional connectivity, we chose to discount changes in absolute signal level and defined the states by their covariance matrix. Inference was run at the group level, such that the state descriptions are defined across subjects. This allowed us to discover dynamic temporal patterns of whole-brain functional interactions along with their occurrence (state time series) and transition probabilities for the duration of the whole resting state fMRI scan. Detailed information about the HMM implementation and the variational Bayes inference can be found in (Vidaurre et al., 2017b,a; Baker et al., 2014). Fractional occupancy was defined as the proportion of time spent on each state and switching rate as the total number of switches from one state to any other. All analyses controlled for age, gender and motion during the resting state fMRI scan.

#### Static functional connectivity

Network modelling was carried out by using the FSLNets toolbox. We calculated the partial temporal correlation between the 15 components’ timeseries creating a 15 ÃŮ 15 matrix of connectivity estimates for each participant. To improve the stability of the estimates of partial correlation coefficients, a small amount of L2 regularisation was applied (Smith et al., 2013a). The connectivity values were converted from Pearson correlation scores into z-statistics with FisherâĂŹs transformation (including an empirical correction for temporal autocorrelation). In order to test the similarity between the correlation matrices (NxN) of each dynamic state and the mean static partial correlation matrix (NxN), we kept the lower diagonal of each matrix, “unwrapped” it to a vector of length 105 ((N*N - N)/2), Fisher’s z transformed its values and calculated the pairwise correlations.

## Acknowledgments

TK was supported by a doctoral studentship of the department of Psychology of the University of York. JS was supported by European Research Council (WANDERINGMINDS - 646927). The authors would like to thank Mladen Sormaz, Charlotte Murphy and Hao-Ting Wang for their contribution to data acquisition.

## References

Baird, B., Smallwood, J., Mrazek, M. D., Kam, J. W., Franklin, M. S., and Schooler, J. W. (2012). Inspired by distraction: mind wandering facilitates creative incubation. Psychological science, 23(10):1117–1122.

Baker, A. P., Brookes, M. J., Rezek, I. A., Smith, S. M., Behrens, T., Smith, P. J. P., and Woolrich, M. (2014). Fast transient networks in spontaneous human brain activity. Elife, 3.

Beaty, R. E., Benedek, M., Wilkins, R. W., Jauk, E., Fink, A., Silvia, P. J., Hodges, D. A., Koschutnig, K., and Neubauer, A. C. (2014). Creativity and the default network: A functional connectivity analysis of the creative brain at rest. Neuropsychologia, 64:92–98.

Beckmann, C. F. and Smith, S. M. (2004). Probabilistic independent component analysis for functional magnetic resonance imaging. IEEE transactions on medical imaging, 23(2):137–152.

Behzadi, Y., Restom, K., Liau, J., and Liu, T. T. (2007). A component based noise correction method (compcor) for bold and perfusion based fmri. Neuroimage, 37(1):90–101.

Cabral, J., Vidaurre, D., Marques, P., Magalhães, R., Moreira, P. S., Soares, J. M., Deco, G., Sousa, N., and Kringelbach, M. L. (2017). Cognitive performance in healthy older adults relates to spontaneous switching between states of functional connectivity during rest. Scientific Reports, 7(1):5135.

Callard, F., Smallwood, J., Golchert, J., and Margulies, D. S. (2013). The era of the wandering mind? twenty-first century research on self-generated mental activity. Frontiers in psychology, 4:891.

Christoff, K., Irving, Z. C., Fox, K. C., Spreng, R. N., and Andrews-Hanna, J. R. (2016). Mind-wandering as spontaneous thought: a dynamic framework. Nature Reviews Neuroscience, 17(11):718.

Craddock, C., Sikka, S., Cheung, B., Khanuja, R., Ghosh, S. S., Yan, C., Li, Q., Lurie, D., Vogelstein, J., Burns, R., et al. (2013). Towards automated analysis of connectomes: The configurable pipeline for the analysis of connectomes (c-pac). Front Neuroinform, 42.

de Silva, P. and Marks, M. (1999). The role of traumatic experiences in the genesis of obsessive–compulsive disorder. Behaviour Research and Therapy, 37(10):941–951.

Engert, V., Smallwood, J., and Singer, T. (2014). Mind your thoughts: associations between self-generated thoughts and stress-induced and baseline levels of cortisol and alpha-amylase. Biological psychology, 103:283–291.

Finn, E. S., Shen, X., Scheinost, D., Rosenberg, M. D., Huang, J., Chun, M. M., Papademetris, X., and Constable, R. T. (2015). Functional connectome fingerprinting: identifying individuals using patterns of brain connectivity. Nature neuroscience, 18(11):1664.

Friston, K. J., Williams, S., Howard, R., Frackowiak, R. S., and Turner, R. (1996). Movement-related effects in fmri time-series. Magnetic resonance in medicine, 35(3):346–355.

Giambra, L. M. (1989). Task-unrelated thought frequency as a function of age: a laboratory study. Psychology and aging, 4(2):136.

Golchert, J., Smallwood, J., Jefferies, E., Seli, P., Huntenburg, J. M., Liem, F., Lauckner, M. E., Oligschläger, S., Bernhardt, B. C., Villringer, A., et al. (2017). Individual variation in intentionality in the mind-wandering state is reflected in the integration of the default-mode, fronto-parietal, and limbic networks. Neuroimage, 146:226–235.

Gonzalez-Castillo, J., Hoy, C. W., Handwerker, D. A., Robinson, M. E., Buchanan, L. C., Saad, Z. S., and Bandettini, P. A. (2015). Tracking ongoing cognition in individuals using brief, whole-brain functional connectivity patterns. Proceedings of the National Academy of Sciences, 112(28):8762–8767.

Greve, D. N. and Fischl, B. (2009). Accurate and robust brain image alignment using boundary-based registration. Neuroimage, 48(1):63–72.

Harris-Warrick, R. M. (2011). Neuromodulation and flexibility in central pattern generator networks. Current opinion in neurobiology, 21(5):685–692.

Hasenkamp, W. and Barsalou, L. W. (2012). Effects of meditation experience on functional connectivity of distributed brain networks. Frontiers in human neuroscience, 6:38.

Hasenkamp, W., Wilson-Mendenhall, C. D., Duncan, E., and Barsalou, L. W. (2012). Mind wandering and attention during focused meditation: a fine-grained temporal analysis of fluctuating cognitive states. Neuroimage, 59(1):750–760.

James, W. (1890). The principles of. Psychology, 2:94.

Karapanagiotidis, T., Bernhardt, B. C., Jefferies, E., and Smallwood, J. (2017). Tracking thoughts: Exploring the neural architecture of mental time travel during mind-wandering. Neuroimage, 147:272–281.

Killingsworth, M. A. and Gilbert, D. T. (2010). A wandering mind is an unhappy mind. Science, 330(6006):932–932.

Konishi, M. and Smallwood, J. (2016). Shadowing the wandering mind: how understanding the mind-wandering state can inform our appreciation of conscious experience. Wiley Interdisciplinary Reviews: Cognitive Science, 7(4):233–246.

Kucyi, A. (2017). Just a thought: how mind-wandering is represented in dynamic brain connectivity. Neuroimage.

Kucyi, A., Esterman, M., Riley, C. S., and Valera, E. M. (2016). Spontaneous default network activity reflects behavioral variability independent of mind-wandering. Proceedings of the National Academy of Sciences, 113(48):13899–13904.

Margulies, D. S., Ghosh, S. S., Goulas, A., Falkiewicz, M., Huntenburg, J. M., Langs, G., Bezgin, G., Eickhoff, S. B., Castellanos, F. X., Petrides, M., et al. (2016). Situating the default-mode network along a principal gradient of macroscale cortical organization. Proceedings of the National Academy of Sciences, 113(44):12574–12579.

Medea, B., Karapanagiotidis, T., Konishi, M., Ottaviani, C., Margulies, D., Bernasconi, A., Bernasconi, N., Bernhardt, B. C., Jefferies, E., and Smallwood, J. (2016). How do we decide what to do? resting-state connectivity patterns and components of self-generated thought linked to the development of more concrete personal goals. Experimental brain research, pages 1–13.

Mooneyham, B. W., Mrazek, M. D., Mrazek, A. J., Mrazek, K. L., Phillips, D. T., and Schooler, J. W. (2017). States of mind: Characterizing the neural bases of focus and mind-wandering through dynamic functional connectivity. Journal of cognitive neuroscience, 29(3):495–506.

Najmi, S., Riemann, B. C., and Wegner, D. M. (2009). Managing unwanted intrusive thoughts in obsessive–compulsive disorder: Relative effectiveness of suppression, focused distraction, and acceptance. Behaviour Research and Therapy, 47(6):494–503.

Poerio, G. L., Totterdell, P., and Miles, E. (2013). Mind-wandering and negative mood: Does one thing really lead to another? Consciousness and cognition, 22(4):1412–1421.

Power, J. D., Mitra, A., Laumann, T. O., Snyder, A. Z., Schlaggar, B. L., and Petersen, S. E. (2014). Methods to detect, characterize, and remove motion artifact in resting state fmri. Neuroimage, 84:320–341.

Quinn, A., Vidaurre, D., Abeysuriya, R., Becker, R., Nobre, A. C., and Woolrich, M. W. (2018). Task-evoked dynamic network analysis through hidden markov modelling. Frontiers in Neuroscience, 12:603.

Ruby, F. J., Smallwood, J., Engen, H., and Singer, T. (2013). How self-generated thought shapes moodâATthe relation between mind-wandering and mood depends on the socio-temporal content of thoughts. PloS one, 8(10):e77554.

Schooler, J. W. (2002). Re-representing consciousness: Dissociations between experience and meta-consciousness. Trends in cognitive sciences, 6(8):339–344.

Seli, P., Kane, M. J., Smallwood, J., Schacter, D. L., Maillet, D., Schooler, J. W., and Smilek, D. (2018). Mind-wandering as a natural kind: A family-resemblances view. Trends in cognitive sciences, 22(6):479–490.

Singer, J. L. and McCraven, V. G. (1961). Some characteristics of adult daydreaming. The Journal of psychology, 51(1):151–164.

Smallwood, J. (2013). Distinguishing how from why the mind wanders: a process–occurrence framework for self-generated mental activity. Psychological bulletin, 139(3):519.

Smallwood, J. and Andrews-Hanna, J. (2013). Not all minds that wander are lost: the importance of a balanced perspective on the mind-wandering state. Frontiers in psychology, 4:441.

Smallwood, J., Karapanagiotidis, T., Ruby, F., Medea, B., de Caso, I., Konishi, M., Wang, H.-T., Hallam, G., Margulies, D. S., and Jefferies, E. (2016). Representing representation: Integration between the temporal lobe and the posterior cingulate influences the content and form of spontaneous thought. PloS one, 11(4):e0152272.

Smallwood, J., Nind, L., and OâAZConnor, R. C. (2009). When is your head at? an exploration of the factors associated with the temporal focus of the wandering mind. Consciousness and cognition, 18(1):118–125.

Smallwood, J. and O’Connor, R. C. (2011). Imprisoned by the past: unhappy moods lead to a retrospective bias to mind wandering. Cognition & emotion, 25(8):1481–1490.

Smallwood, J. and Schooler, J. W. (2006). The restless mind. Psychological bulletin, 132(6):946.

Smallwood, J. and Schooler, J. W. (2015). The science of mind wandering: empirically navigating the stream of consciousness. Annual review of psychology, 66:487–518.

Smith, J. M. and Alloy, L. B. (2009). A roadmap to rumination: A review of the definition, assessment, and conceptualization of this multifaceted construct. Clinical psychology review, 29(2):116–128.

Smith, S. M., Beckmann, C. F., Andersson, J., Auerbach, E. J., Bijsterbosch, J., Douaud, G., Duff, E., Feinberg, D. A., Griffanti, L., Harms, M. P., et al. (2013a). Resting-state fmri in the human connectome project. Neuroimage, 80:144–168.

Smith, S. M., Hyvärinen, A., Varoquaux, G., Miller, K. L., and Beckmann, C. F. (2014). Group-pca for very large fmri datasets. Neuroimage, 101:738–749.

Smith, S. M., Nichols, T. E., Vidaurre, D., Winkler, A. M., Behrens, T. E., Glasser, M. F., Ugurbil, K., Barch, D. M., Van Essen, D. C., and Miller, K. L. (2015). A positive-negative mode of population covariation links brain connectivity, demographics and behavior. Nature neuroscience, 18(11):1565.

Smith, S. M., Vidaurre, D., Beckmann, C. F., Glasser, M. F., Jenkinson, M., Miller, K. L., Nichols, T. E., Robinson, E. C., Salimi-Khorshidi, G., Woolrich, M. W., et al. (2013b). Functional connectomics from resting-state fmri. Trends in cognitive sciences, 17(12):666–682.

Sormaz, M., Murphy, C., Wang, H.-t., Hymers, M., Karapanagiotidis, T., Poerio, G., Margulies, D. S., Jefferies, E., and Smallwood, J. (2018). Default mode network can support the level of detail in experience during active task states. Proceedings of the National Academy of Sciences.

Stawarczyk, D., Majerus, S., Maj, M., Van der Linden, M., and D’Argembeau, A. (2011). Mind-wandering: phenomenology and function as assessed with a novel experience sampling method. Acta psychologica, 136(3):370–381.

Vatansever, D., Menon, D. K., and Stamatakis, E. A. (2017). Default mode contributions to automated information processing. Proceedings of the National Academy of Sciences, 114(48):12821–12826.

Vidaurre, D., Abeysuriya, R., Becker, R., Quinn, A. J., Alfaro-Almagro, F., Smith, S. M., and Woolrich, M. W. (2017a). Discovering dynamic brain networks from big data in rest and task. Neuroimage.

Vidaurre, D., Quinn, A. J., Baker, A. P., Dupret, D., Tejero-Cantero, A., and Woolrich, M. W. (2016). Spectrally resolved fast transient brain states in electrophysiological data. Neuroimage, 126:81–95.

Vidaurre, D., Smith, S. M., and Woolrich, M. W. (2017b). Brain network dynamics are hierarchically organized in time. Proceedings of the National Academy of Sciences, 114(48):12827–12832.

Vidaurre, D., Woolrich, M. W., Winkler, A. M., Karapanagiotidis, T., Smallwood, J., and Nichols, T. E. (2018). Stable between-subject statistical inference from unstable within-subject functional connectivity estimates. Human Brain Mapping.

Watkins, E. R. (2008). Constructive and unconstructive repetitive thought. Psychological bulletin, 134(2):163.

